# Primary Cilia Dysfunction in Brown Fat Results in Fatal Thermogenesis Failure in Neonatal Mice

**DOI:** 10.1101/2025.11.12.687971

**Authors:** Yong Min Kim, Hyeseung Lee, Joo Heon Kim, Jaeku Kang, Seung Yun Han, Jin Sun Park, Hwan-Woo Park, Jongdae Shin, Jong-seok Kim, Hyung Seo Park, Junguee Lee

## Abstract

Primary cilium, microtubule-based sensory organelle that has emerged as a central player in coordinating numerous signaling pathways. Although primary cilia are known to regulate cellular signaling and energy metabolism, the relationship between these roles in the context of ciliopathies and brown adipose tissue (BAT) dysfunction remains poorly understood. To elucidate the role of primary cilia in BAT, we generated *Ucp1-Cre;Ift88^flox/flox^* mice with BAT-specific ciliary loss. There were no significant differences in body size, weight, or BAT weight relative to body weight between P0 *Ucp1-Cre;Ift88^flox/flox^* and P0 *Ucp1-Cre;Ift88^+/+^* pups. Embryos examined at E14.5 and E18.5 also showed no discernible differences in size, morphology, or histology. However, P0 *Ucp1-Cre;Ift88^flox/flox^* pups exhibited neonatal lethality caused by defective thermogenesis, despite preserved Ucp1 expression. These mice displayed markedly reduced ketone body levels in both BAT and serum, accompanied by downregulation of *Hmgcs2*, a key enzyme in ketogenesis. Loss of primary cilia in BAT suppressed ketogenesis and increased ROS production through HMGCS2 downregulation, ultimately impairing non-shivering thermogenesis. Remarkably, neonatal lethality of *Ucp1-Cre;Ift88^flox/flox^* pups was completely rescued by thermoneutral housing or β-hydroxybutyrate supplementation. Our findings identified a previously unrecognized mechanism by which primary cilia regulate non-shivering, UCP1-independent thermogenesis via ketogenesis.

## Introduction

The primary cilium is a solitary microtubule-organized sensory organelle that has emerged as a key regulator of numerous signaling pathways. Present on nearly all mammalian cells, the primary cilium functions as a cellular antenna that detects extracellular signals and transduces them into intracellular responses^1^. Diseases characterized by dysfunctional primary cilia, known as ciliopathies, exhibit diverse clinical features, including obesity and diabetes^2^. Bardet-Biedl syndrome arises from impaired primary cilia caused by mutations in individual members of the *Bbs* gene family^3^. Recently, Gohlke *et al*. demonstrated that the loss of *Bbs4* impairs thermogenic potential, leading to cold intolerance^4^. This impairment was linked to defective lipid mobilization in adipose tissue and liver, as well as altered fatty acid metabolism^5^. Their study highlighted the functional link between primary cilia and thermogenesis, revealing how the BBSome influences energy metabolism and lipid utilization in brown and white adipocytes. By studying whole-body *Bbs4* knockout mice, they underscored the connection between primary cilia and thermogenesis.

BAT plays a crucial role in thermogenesis and energy expenditure in mammals^5^. Unlike white adipose tissue (WAT), which stores energy as lipids, BAT oxidizes lipids to generate heat^6^. This thermogenic process is enabled by mitochondrial uncoupling protein 1 (UCP1)^6^. In neonatal mice, the function of BAT is particularly crucial. BAT reaches its maximum relative weight at birth and gradually decreases during the first 4 to 5 days of postnatal days^7^. Neonates, with limited energy reserves and a high surface-to-volume ratio, are highly vulnerable to heat loss and hypothermia^8^. Unlike adults, they cannot effectively generate heat through shivering and therefore depend heavily on non-shivering thermogenesis provided by BAT for survival^9^. BAT achieves this by oxidizing fatty acids and glucose to generate heat, a process facilitated by UCP1^10^. The initiation of thermogenesis in response to cold exposure involves the release of norepinephrine by the sympathetic nervous system^5^. This neurotransmitter binds to the β3-adrenergic receptors on brown adipocytes, triggering the breakdown of stored lipids into free fatty acids, which subsequently activate UCP1 to promote heat production^5^. Beyond thermogenesis, BAT significantly contributes to overall metabolism in neonatal mice^11^. The high metabolic activity of BAT not only supports heat generation but also regulates body weight and energy balance. Although primary cilia are recognized for their roles in cellular signaling and energy metabolism, their contribution to BAT dysfunction in the context of ciliopathies remains poorly defined.

This study examines the role of primary cilia in BAT regulation in neonatal mice. We disrupted the Ift88 gene in BAT to induce BAT-specific primary cilia dysfunction. Neonatal mice with BAT-specific Ift88 deletion exhibited early postnatal lethality caused by compromised thermoregulation. Loss of primary cilia or disruption of ciliogenesis in BAT led to impaired ketogenesis and increased reactive oxygen species (ROS) levels. These defects were attributed to decreased Hmgcs2, a key enzyme in ketogenesis, resulting in the failure of non-shivering thermogenesis. Overall, these findings underscore the significant role for primary cilia in BAT thermogenesis and energy regulation, providing novel insights into the pathophysiology of ciliopathies and suggesting potential therapeutic strategies.

## Materials and Methods

### Generation of BAT-Specific Ift88 knockdown mice

To disrupt primary cilia in brown fat, *Ucp1-Cre* males (obtained from Prof. Shong M) were crossed with *Ift88^flox/flox^* female mice (obtained from Prof. Kim J, KAIST). Offspring carrying *Ucp1-Cre;Ift88^flox/+^* mice exhibited reduced viability of the homozygous floxed genotype, as *Ucp1-Cre;Ift88^flox/flox^* pups died shortly after birth. The genetic inheritance at embryonic day 14.5 (E14.5) and E18.5 embryos was as expected by the Hardy–Weinberg equilibrium. For inducible deletion, *Ucp1-CreERT2* males (purchased from Prof. Shong M) were bred with *Ift88^flox/flox^* female mice. Animal experiments received prior approval by the Institutional Animal Care and Use Committee (IACUC) of Konyang University (KYU, approval ID: KY-IACUC-24-10-C-01), and the IACUC of KYU follows the rules of Basel Declaration. KYU is an Association for Assessment and Accreditation of Laboratory Animal Care International (AAALAC International)-accredited facility and abides by the Institute of Laboratory Animal Resources (ILAR) guide.

### Cell lines and cell culture

Murine *i*BPA cell lines (Merck, cat#SCC255, USA) were maintained in Dulbecco’s modified Eagle’s medium (DMEM) supplemented with 10% fetal calf serum and 10,000 U/ml Penicillin-Streptomycin at 37°C in a 5% CO₂ atmosphere. Once cells reached confluence, they were incubated for 48 hrs in a differentiation-inducing medium containing 10 μg/ml insulin and 2.5 μM dexamethasone. Following induction, the cells were further cultured in a maturation medium supplemented with 10 μg/ml insulin for an additional 5 to 7 days.

### In vivo Infrared Thermography

BAT temperature was noninvasively estimated by measuring the skin surface temperature on interscapular BAT with a high-sensitivity infrared thermography camera (T650sc, emissivity of 0.98, FLiR Systems). Neonatal mice were housed at temperature of 26°C in open-top cages, each positioned under a thermal camera. Static dorsal thermograms were acquired within 12 hrs after birth using an infrared camera mounted on top of a box placed above the mouse on the cage grid, with a focal length of 20 cm. Images were analyzed in FLiR Research IR software using the rainbow high-contrast color palette.

### RNA sequencing analysis

BATs extracted from three P0 mice per group (*Ucp1-Cre;Ift88^flox/flox^, Ucp1-Cre;Ift88^+/+^* mice) were used for RNA sequencing. Because gender distinction of neonatal pup is difficult, male and female mice were used in the current study. Mice were sacrificed and BAT isolated. Extracted BATs were preserved in liquid nitrogen and transported to Macrogen Inc. (South Korea) for RNA sequencing analysis. A TruSeq Stranded Total RNA LT Sample Prep Kit (Gold) was used as a library kit, and the NovaSeq 6000 S4 Reagent Kit was used as a reagent. DEG analysis was conducted on three comparison combinations using FPKM, and 1473 genes were extracted from at least one comparison combination that met |fold change| ≥ 2 and independent t-test raw *p*-value < 0.05.

### Total RNA isolation and RT-qPCR

Total RNA was extracted from mouse BAT and cultured cells using TRIzol™ Plus RNA Purification Kit (Thermo Fisher Scientific Inc). Complementary DNA (cDNA) was synthesized from the extracted RNA using M-MLV Reverse Transcriptase and oligo-dT primers (Invitrogen). Reverse transcription quantitative PCR (RT-qPCR) was conducted on an Applied Biosystems 7500 Real-Time PCR System (Thermo Fisher Scientific Inc.) with the QuantiTect SYBR Green PCR Master Mix (QIAGEN). Reactions were performed in triplicate, with the following amplification protocol: 10 min at 95°C for enzyme activation, 10 sec at 95°C for denaturation, followed by 40 cycles of 30 sec at 60°C for annealing and 30 sec at 72°C for extension. Gene expression was quantified using the delta cycle threshold (Ct) method, with normalization to *Gapdh* expression, and the data were analyzed using Life Technologies 7500 software (Foster City, CA, USA).

### Immunoblot analysis

Mouse BATs were homogenized using RIPA lysis buffer supplemented with a protease inhibitor cocktail. Protein concentrations were determined using the Bradford assay and samples were prepared by adding SDS sample buffer. The samples were denatured by heating at 95°C for 5 min, applied to a polyacrylamide gel for electrophoresis. Following electrophoresis, proteins were transferred to a nitrocellulose membrane using the wet transfer method. The membrane was then incubated with blocking buffer for 30 min and subsequently incubated overnight at 4°C with the corresponding primary antibodies specific for IFT88 (Proteintech, 1:1000), HMGCS2 (Abclonal, 1:1000), α-Tubulin (Sigma-Aldrich, 1:1000) and β-Actin (Abcam, 1:1000). Following several washes to remove excess primary antibody, the membrane was exposed with a secondary antibody for 2 hrs at room temperature. The membrane was washed three times for 10 min each with TBS/T, then visualized using a chemiluminescent reagent kit (Immobilon Western Chemiluminescent HRP Substrate, Merck Millipore). α-Tubulin and β-Actin were used as a loading control. Band densities on the immunoblots were quantified using ImageJ gel analysis.

### Immunofluorescence

Paraffin-embedded tissue sections (4 μm) were incubated at 56°C for 3 hrs, then deparaffinized in xylene and rehydrated through graded ethanol. Antigen retrieval was performed in 0.01 M citrate buffer (pH 6.0) using an autoclave at 121°C for 25 min. Slides were air-dried for 30 min and rinsed with 1× PBS. Tissue slice sections were stained using the BenchMark ULTRA PLUS system (Roche, Switzerland). Primary antibody was anti-ARL13B (Proteintech, 1:400), polyglutamylation modification (GT335, AdipoGen Life Sciences, 1:400), γ-Tubulin (Sigma-Aldrich, 1:400), HMGCS2 (Abclonal, 1:500), and Adenylate Cyclase 3 (Invitrogen, 1:100) were used. Goat anti-mouse and goat anti-rabbit secondary antibodies conjugated to Alexa Fluor 488 or 568 (Invitrogen/Life Technologies, 1:400) were used for indirect IF. The stained slides were observed under the LSM 700 Laser Scanning Microscope (Carl Zeiss).

### Immunohistochemistry

Paraffin-embedded 4 μm-thick tissue slices were placed in an oven and incubated at 56°C for 3 hrs. Immunohistochemistry (IHC) was performed using the BenchMark ULTRA PLUS system (Roche Diagnostics, United States). The primary antibody was anti-UCP1 (Abcam, 1:100), anti-Perilipin (Abcam, 1:100), and recombinant anti-HMGCS2 antibody (Abcam, 1:100).

### β3-adrenergic stimulation

Mice were housed under thermoneutral condition (30°C) and intraperitoneally injected with β3-adrenergic agonist CL-316243 hydrate (Sigma-Aldrich, 0.5 ㎍/g body weight/day). Rectal temperature was measured at 3 hrs after injection. To assess of the thermogenic response, BAT was harvested 3 hrs post-injection for qPCR and western blot analysis of Ucp1 and mitochondrial gene expression.

### Measurement of mitochondrial respiration and glycolysis in BAT

Oxygen consumption rate (OCR) and extracellular acidification rate (ECAR) were measured using the Seahorse XF24 Analyzer (Agilent Technologies), following a modified protocol optimized for BAT slices^12^. Briefly, interscapular BAT was freshly isolated and cut into approximately 1 mm³ tissue fragments under sterile conditions. Each tissue slice was carefully placed into a well of an XF24 Islet Capture Microplate (Agilent Technologies) using the manufacturer-supplied islet capture screens to immobilize the tissue. Each well contained 200 µL of assay medium pre-warmed to 37 °C. The assay medium consisted of unbuffered DMEM (pH 7.4) supplemented with 25 mM glucose, 1 mM pyruvate, and 2 mM glutamine. Plates were incubated at 37°C in a non-CO₂ incubator for 1 hr to allow temperature and pH equilibration.

On the day of the assay, the XF sensor cartridge was hydrated overnight in XF Calibrant at 37°C and calibrated according to the manufacturer’s instructions. Following equilibration, an additional 300 µL of pre-warmed assay medium was added to each well, bringing the total volume to 500 µL.

OCR was recorded under basal conditions and after sequential injections of mitochondrial inhibitors using the XF24 Analyzer: oligomycin (2 µM, ATP synthase inhibitor), carbonyl cyanide-p-trifluoromethoxyphenylhydrazone (FCCP; 10 µM, a mitochondrial uncoupler), and rotenone (1 µM, a complex I inhibitor). Three measurement cycles (mix, wait, and measure) were performed after each injection.

Data were analyzed using Seahorse XF24 software (Agilent), and OCR values were normalized to total protein content per well. Following the assay, tissues were lysed, and protein concentrations were quantified using a BCA protein assay (Thermo Scientific) to ensure equal tissue loading across wells.

### BAT membrane phospholipid analysis

BATs were extracted from six P0 mice per sample immediately after birth. Whole-tissue membrane lipids were then extracted from about 15 mg of BAT (6 mice/sample). The extracted BATs were sent to NICEM Chromatography center (College of Agriculture and Life Sciences, Seoul National University, South Korea) for Gas chromatography analysis (Agilent 7890A).

### Determination of Ketone bodies in BAT

BATs extracted from six P0 mice per sample. Extracted BATs were sent to Chungnam National Univ. Center for Research Facilities (South Korea) for Gas chromatography mass spectrometry (GC-MS)(Perkinelmer. MS 2400 SQ).

### Ketone body supplementation

Within 1 hr of birth, all P0 *Ucp1-Cre;Ift88* floxed pups received intraperitoneal injections of β-OHB (DL-β-Hydroxybutyryl coenzyme A lithium salt, Sigma-Aldrich) at 0.1 mg/g, then twice daily for 2 successive days. These mice were genotyped by PCR analysis of tail DNA at 2 weeks of age.

### Knockdown of Ift88 using RNA interference

Cells were plated in 6-well or 12-well cell plate and allowed to adhere overnight. Cells were transfected with 30 nM Ift88 (Assay ID 186729, Thermo Fisher Scientific Inc.) in Opti-MEM I medium using Lipofectamine RNAiMAX transfection reagent (Invitrogen) following the manufacturer’s instructions. A non-targeting siRNA with no sequence similarity to the human genome, supplied by Invitrogen, was used as a negative control. All experiments were conducted in triplicate and independently repeated at least three times. Gene knockdown efficiency was assessed by RT-qPCR 48 hrs post-transfection. Additionally, Immunofluorescence (IF) imaging confirmed a reduction in the number of primary cilia in the experimental group compared to the control.

### Flow cytometric analysis for ROS

Cells transfected with si*Kif3a* or si*Ift88* were harvested and washed twice with Hanks’ Balanced Salt Solution (HBSS). To analyze ROS product, the cells were resuspended in HBSS and then incubated with 5 µM of 5-(and-6)-carboxy-2’,7’-dichlorodihydro fluorescein diacetate (Carboxy-DCFDA; Sigma-Aldrich, St Louis, MO, USA) in HB2S for 15 min at 37°C in darkness. After incubated then washed 2 times with HBSS for the fluorescent adduct (DCF signal) of the collected cells was measured using FACS Canto-II flow cytometer (BD Biosciences, USA).

### Intracellular ATP assay

Intracellular ATP concentration was determined by an ATP fluorometric assay kit (Abcam), based on the procedure recommended by the manufacturer.

### Statistical analysis

Data are presented as the mean ± standard error (SEM). Differences between two groups were assessed using an unpaired Student’s t-test, with a *p*-value less than 0.05 considered indicative of a significant difference.

## Results

### Generation and characterization of Ucp1-Cre;Ift88 ^flox/flox^ mice

To investigate the function of primary cilia in BAT, *Ucp1-Cre;Ift88* floxed mice were generated using the *Cre/loxP* recombination system. Genotypes were confirmed by PCR analysis of tail genomic DNA (Figure 1A), and *Ift88* knockdown was verified by immunoblotting (Figure 1B). Because *Ift88* is critical for ciliary assembly and maintenance, we examined whether primary cilia were lost in BAT of *Ucp1-Cre;Ift88^flox/flox^* mice using IHC. Given that *Ucp1-Cre* is activated around embryonic day (E) 16.5 and mature brown adipocytes typically lack primary cilia, BAT was harvested immediately before birth. Primary cilia were readily detected in BAT of *Ucp1-Cre;Ift8^+/+^* mice but were absent in *Ucp1-Cre;Ift88^flox/flox^* littermates (Figure 1C).

**Figure 1.**
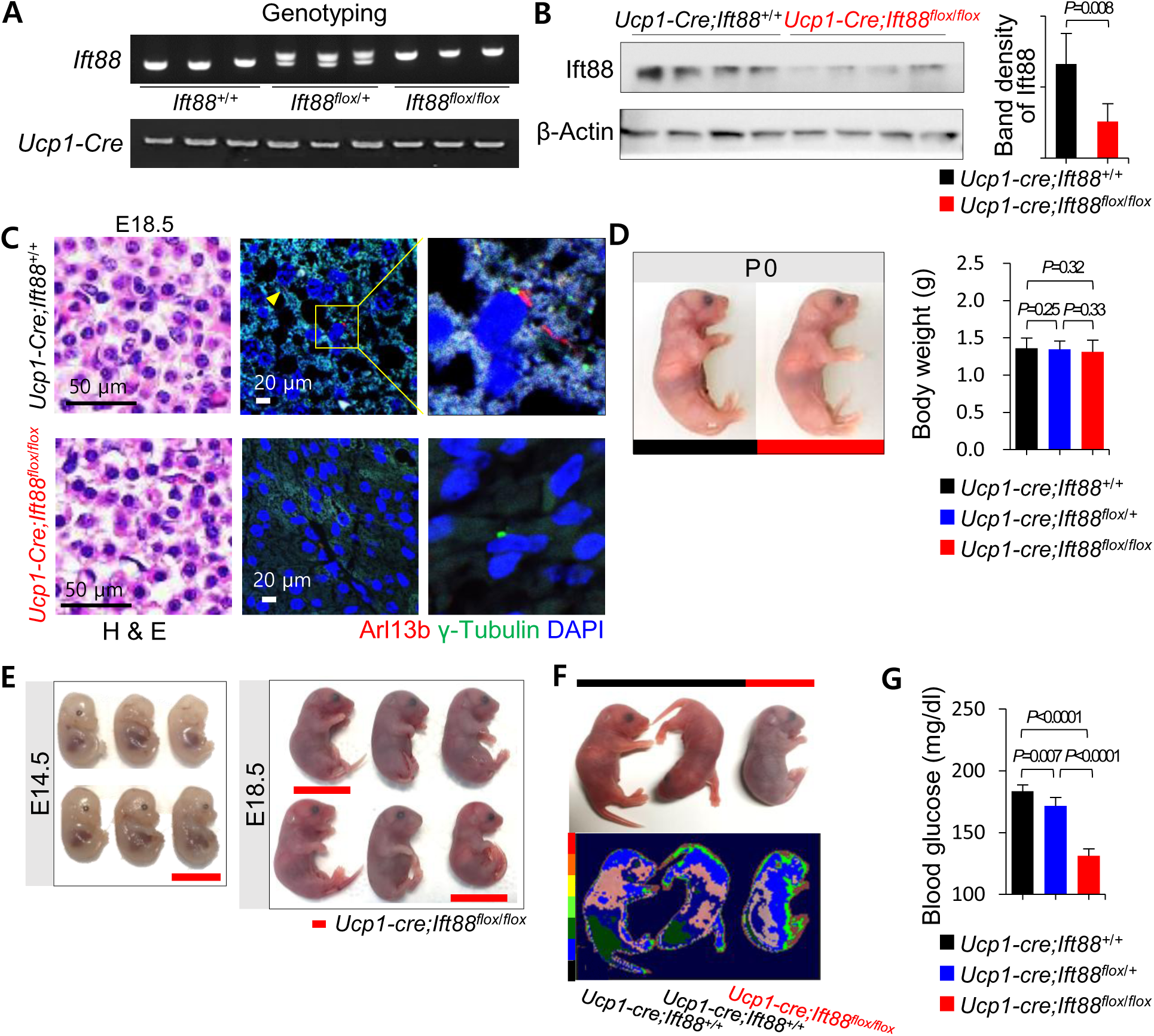
Generation and validation of *Ucp1-Cre;Ift88* floxed mice. (A) The *Ift88^+/+^*, *Ift88^+/-^*, and *Ift88^-/-^*genotypes are analyzed by PCR using mouse tail DNA. (B) Immunoblot analysis shows a reduced level of IFT88 in the BAT of *Ucp1-Cre;Ift88^flox/flox^* mice compared to *Ucp1-Cre;Ift88^+/+^* mice (*p=0.008*). β-Actin is used as the loading control. (C) The presence of primary cilia is confirmed by IF with Arl13b (red) for the axoneme and γ-Tubulin (green) for the basal body. In E18.5 *Ucp1-Cre; Ift88^flox/flox^* BAT, primary cilia are rarely detected. (D) The body weight of P0 *Ucp1-Cre;Ift88^+/+^*, *Ucp1-Cre;Ift88^flox/+^*, and *Ucp1-Cre; Ift88^flox/flox^* pups are 1.36 ± 0.13 g, 1.35 ± 0.11 g, and 1.31 ± 0.15 g, respectively. (E) The gross appearance of E14.5 or E18.5 embryos among the groups. (F) Infrared thermography specifically detects BAT thermogenesis in wild type and *Ucp1-Cre; Ift88^flox/flox^* pups at 12 hrs after birth. Representative infrared images demonstrate impaired BAT thermogenesis in *Ucp1-Cre; Ift88^flox/flox^* pups. (G) Blood glucose levels in P0 *Ucp1-Cre;Ift88^+/+^*, *Ucp1-Cre;Ift88^flox/+^*, and *Ucp1-Cre; Ift88^flox/flox^* pups are 183.6 ± 5.13 mg/dl, 171.6 ± 6.88 mg/dl, and 131.2 ± 5.54 mg/dl, respectively.

Unexpectedly, *Ucp1-Cre;Ift88^flox/flox^* pups exhibited postnatal mortality within 24 hrs after birth. No significant differences in body size and weight were observed between postnatal day zero (P0) *Ucp1-Cre;Ift88^flox/flox^* and *Ucp1-Cre;Ift88^+/+^* pups (Figure 1D). To further assess developmental differences, embryos were examined at E14.5 and E18.5, but no gross morphological abnormalities were detected (Figure 1E). At P0, *Ucp1-Cre;Ift88^+/+^* pups displayed active shivering, whereas the *Ucp1-Cre;Ift88^flox/flox^* newborn pups remained largely immobile at an ambient temperature of 25°C (Supplementary video 1). Infrared thermography, a non-invasive technique for specifically detecting BAT thermogenesis *in vivo*^13^, revealed markedly impaired BAT thermogenesis in the *Ucp1-Cre;Ift88^flox/flox^* pups (Figure 1F).

Consistent with these findings, blood glucose levels were significantly lower in P0 *Ucp1-Cre;Ift88^flox/flox^* mice compared to *Ucp1-Cre;Ift88^+/+^* littermates (Figure 1G). Although hypoglycemia may contribute to the observed hypothermia, the precise causal relationship remains unclear.

### Histological characteristics of BAT in Ucp1-Cre;Ift88 ^flox/flox^ mice

To investigate the histological differences in BAT, *Ucp1-Cre;Ift88^flox/flox^* and *Ucp1-Cre;Ift88^+/+^* pups were sacrificed immediately after birth. Although gross morphology appeared similar between P0 *Ucp1-Cre;Ift88^flox/flox^* and *Ucp1-Cre;Ift88^+/+^* mice, BAT size was visibly smaller in *Ucp1-Cre;Ift88^flox/flox^* mice (Figure 2A). Moreover, quantitative analysis confirmed a significant reduction in BAT weight relative to body weight in *Ucp1-Cre;Ift88^flox/flox^* mice compared to control littermates (Figure 2A).

**Figure 2.**
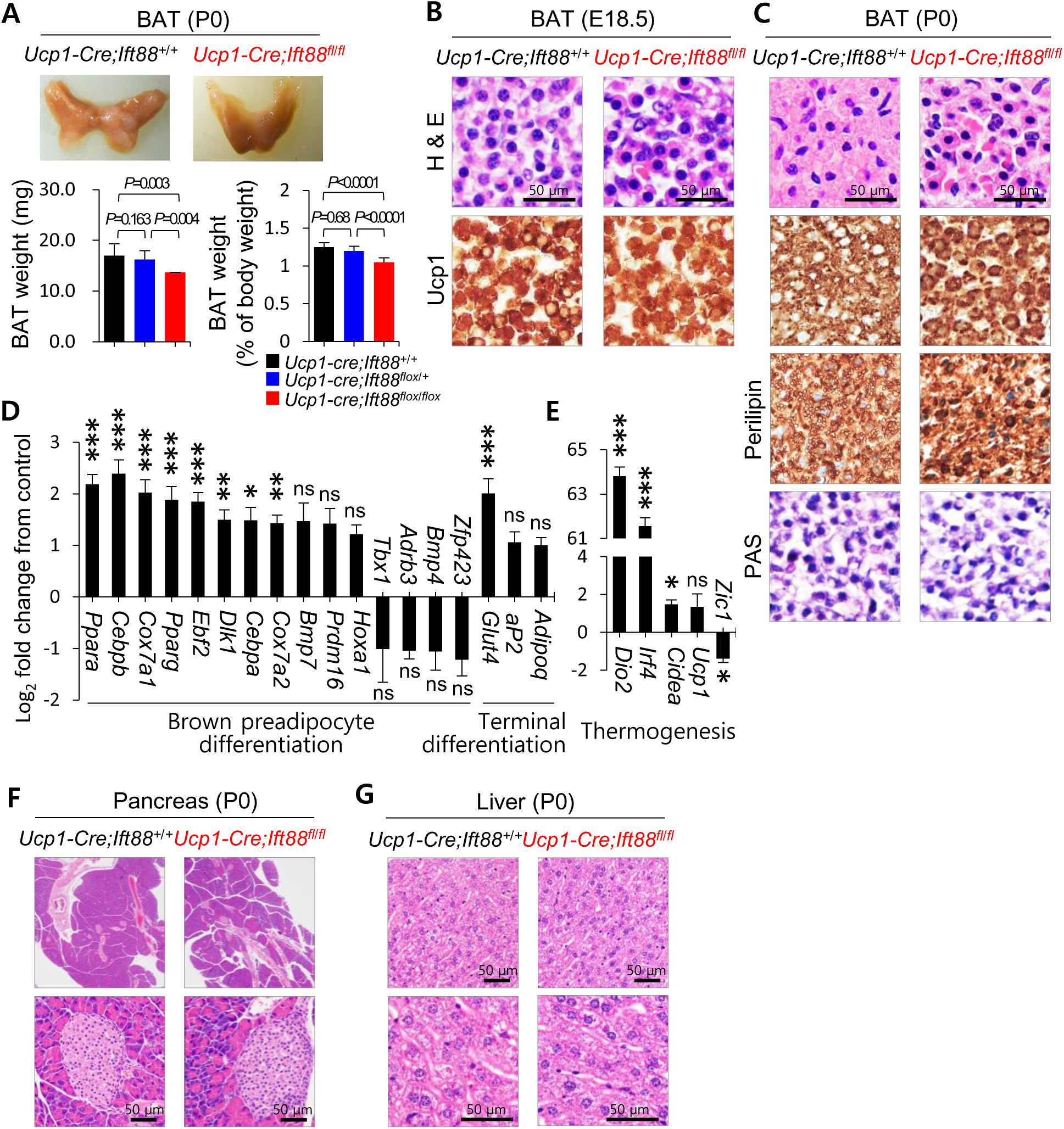
Histology characteristics of BAT in *Ucp1-Cre; Ift88^flox/flox^* neonates. (A) Gross morphology of BAT in P0 *Ucp1-Cre; Ift88^flox/flox^* and P0 *Ucp1-Cre;Ift88^+/+^* mice. The BAT weight of P0 *Ucp1-Cre;Ift88^+/+^*, *Ucp1-Cre;Ift88^flox/+^*, and *Ucp1-Cre; Ift88^flox/flox^* pups are 16.99 ± 2.32 mg, 16.12 ± 1.71 mg, and 13.70 ± 0.02 mg, respectively. The BAT weight of *Ucp1-Cre; Ift88^flox/flox^* pups is significantly lower. (B) At E18.5, BAT histology and Ucp1 expression were comparable between *Ucp1-Cre; Ift88^flox/flox^* and *Ucp1-Cre;Ift88^+/+^* embryos. (C) H&E staining revealed fewer intracytoplasmic lipid vacuoles in BAT of P0 *Ucp1-Cre; Ift88^flox/flox^* mice compared to P0 *Ucp1-Cre;Ift88^+/+^* mice. Intracytoplasmic lipid droplets are demonstrated using perilipin staining. There is no difference of Ucp1 or PAS expression between BAT of P0 *Ucp1-Cre; Ift88^flox/flox^* and BAT of *Ucp1-Cre;Ift88^+/+^* mice. (D, E) BAT from *Ucp1-Cre; Ift88^flox/flox^* pups exhibit similar or higher expressions of genes related to development, differentiation, and thermogenesis compared to *Ucp1-Cre;Ift88^+/+^* BAT. (F, G) Histological features of the pancreas and liver in P0 *Ucp1-Cre; Ift88^flox/flox^* mice are indistinguishable from those of P0 *Ucp1-Cre;Ift88^+/+^* mice.

At E18.5, BAT from both groups showed comparable microscopic finding or Ucp1 expression (Figure 2B). However, P0 *Ucp1-Cre;Ift88^flox/flox^* BAT displayed fewer intracytoplasmic lipid vacuoles compared to that of P0 *Ucp1-Cre;Ift88^+/+^* mice, as shown by H&E staining (Figure 2C). Immunostaining for perilipin, a lipid droplet-associated protein, confirmed the presence of intracytoplasmic lipid droplets in P0 *Ucp1-Cre;Ift88^+/+^* BAT, whereas lipid droplet was markedly reduced in P0 *Ucp1-Cre;Ift88^flox/flox^* BAT (Figure 2C). Ucp1 expression, however, was comparable between groups (Figure 2C).

In normal BAT, absorbed glucose is converted into triglycerides (TGs) through de novo lipogenesis and stored in lipid droplets, providing a critical fuel source for thermogenesis^14^. Therefore, the paucity of lipid droplets in P0 *Ucp1-Cre;Ift88^flox/flox^* BAT suggests possible defects in BAT maturation, impaired lipid synthesis, inefficient conversion of glucose into lipids, or rapid lipid consumption without adequate replenishment.

To elucidate the mechanism underlying the premature mortality of mice lacking primary cilia in BAT, we performed RNA-sequencing analysis (RNA-seq) on BAT samples from *Ucp1-Cre;Ift88* floxed mice (Supplementary dataset). Integrating transcriptomic and histological data revealed alterations in metabolic and thermogenic pathways that may account for the impaired BAT function and neonatal lethality in *Ucp1-Cre;Ift88^flox/flox^* mice.

RNA-seq analysis revealed that genes associated with development and differentiation were expressed at comparable or higher levels in BAT of P0 *Ucp1-Cre;Ift88^flox/flox^* mice relative to controls (Figure 2D). Notably, transcription factors critical for brown adipocyte differentiation, including *Pparg*, *Cebpa*, *Cebpb*, and *Ebf2*, were significantly upregulated in P0 *Ucp1-Cre;Ift88^flox/flox^* BAT (Figure 2D). Likewise, maturation markers such as *Ucp1*, *Cidea*, and *Pgc1a*were comparable and in some cases upregulated (Figure 2D-2E), indicating that BAT maturation was not delayed.

Next, PAS staining was performed to investigate whether glucose was accumulating within cells instead of being converted into lipids. Neither P0 *Ucp1-Cre;Ift88^+/+^* nor *Ucp1-Cre;Ift88^flox/flox^* BAT showed PAS positivity, suggesting that glucose was consumed rather than stored as glycogen (Figure 2C). RNA-seq further revealed upregulation of triglyceride and fatty acid synthesis pathways in P0 *Ucp1-Cre;Ift88^flox/flox^* BAT, indicating that lipid biosynthesis was not impaired.

RNA-seq also identified *GLUT4* as a highly upregulated gene in P0 *Ucp1-Cre;Ift88^flox/flox^* BAT compared to controls (Figure 2D). Consistent with this finding, P0 *Ucp1-Cre;Ift88^flox/flox^* pups exhibited hypoglycemia (Figure 1G). Because glucose supply is limited during the early postnatal day, excessive glucose uptake by P0 *Ucp1-Cre;Ift88^flox/flox^* BAT in these pups is likely to contributes to systemic hypoglycemia. Importantly, histological examination of the pancreas and liver revealed no abnormalities in P0 *Ucp1-Cre;Ift88^flox/flox^* pups compared to controls (Figure 2F-2G). Therefore, the systemic hypoglycemia and hypothermia observed in P0 *Ucp1-Cre;Ift88^flox/flox^* mice are most likely attributable to dysregulated glucose metabolism in BAT, rather than intrinsic defects in the pancreas or liver.

Collectively, these results suggest that in P0 *Ucp1-Cre;Ift88^flox/flox^* BAT, glucose is rapidly utilized rather than stored in lipid droplets.

### Transcriptomic reprogramming associated with hypothermia in P0 Ucp1-Cre;Ift88^flox/flox^ mice

We compared the transcriptomic profiles of BAT from *Ucp1-Cre;Ift88^flox/flox^* and *Ucp1-Cre;Ift88^+/+^* mice. Hierarchical clustering heatmap of differentially expressed genes (DEGs) with a ≥ 2-fold change and *p*-value < 0.05 demonstrated distinct clustering of the two groups, indicating divergent molecular signatures (Figure 3A). Gene set enrichment analysis (GSEA) revealed enrichment of pathways related to oxidative phosphorylation, G2/M checkpoint, E2F targets, cholesterol homeostasis, mTORC1 signaling, glycolysis, hypoxia, TNFα signaling via NF-κB, mitotic spindle, adipogenesis, IL2-STAT5 signaling, fatty acid metabolism, and MYC targets V1 in BAT of *Ucp1-Cre;Ift88^flox/flox^* mice (Figure 3B).

**Figure 3.**
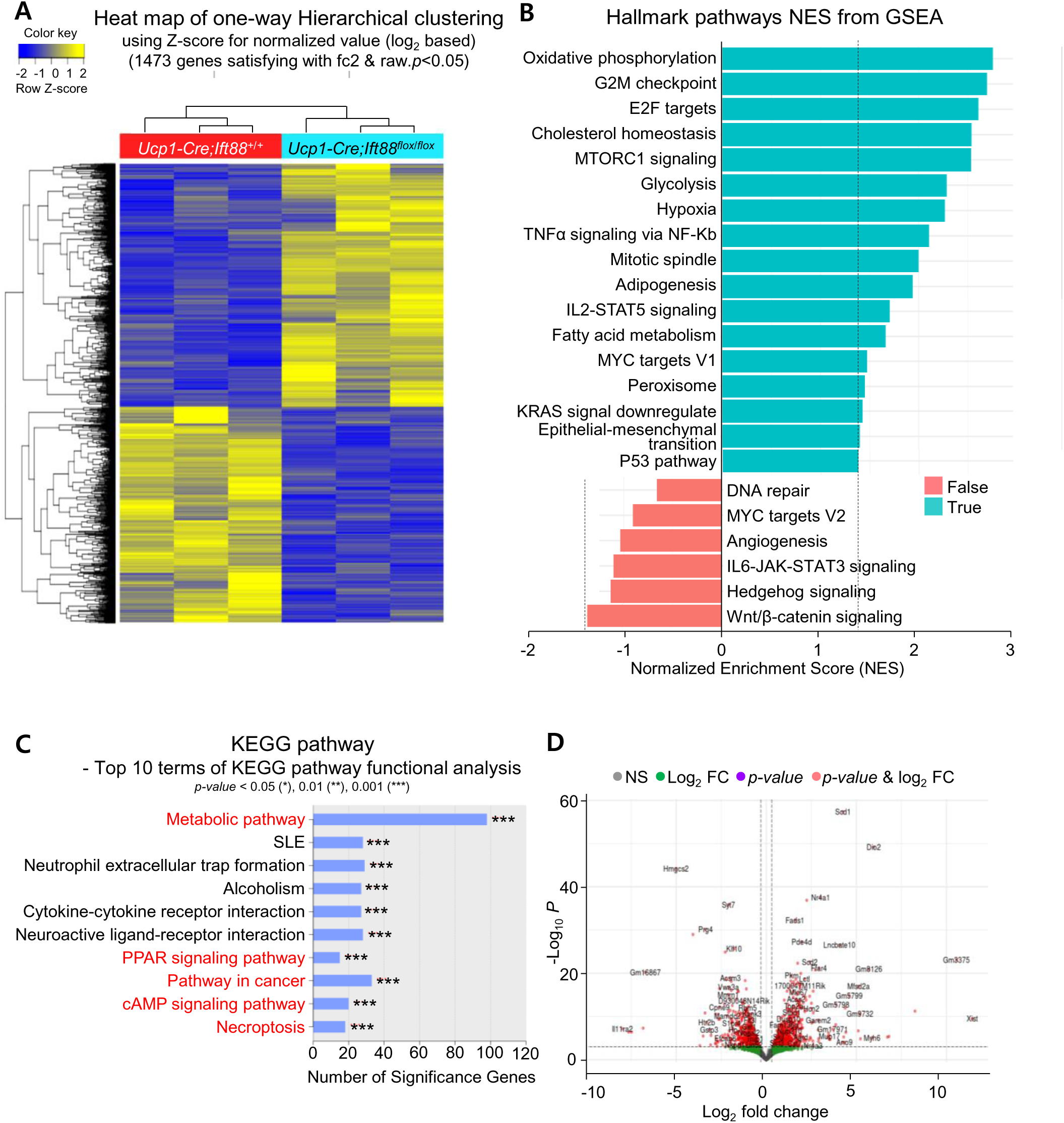
Distinct gene expressions profiles in BAT of P0 *Ucp1-Cre; Ift88^flox/flox^* mice revealed by RNA-sequencing analysis. (A) Hierarchical clustering heatmap of differentially expressed genes in BAT from *Ucp1-Cre; Ift88^flox/flox^* versus *Ucp1-Cre;Ift88^+/+^* pups, satisfying the arbitrary criterion of a ≥ 2-fold change in expression with a *p*-value < 0.05. (B) Waterfall plot of Hallmark gene set enrichment analysis (GSEA) signatures ranked by normalized enrichment score (NES). Positive NES indicates enriched signatures, whereas negative NES indicates depleted signatures. Dotted line representsthe significance cutoff (*p-*value < 0.05). (C) Top 10 KEGG pathway terms identified by functional enrichment analysis results. (D) Volcano plot of RNA-seq data showing 316 upregulated and 208 downregulated genes.

To further characterize DEGs with ciliary loss, we conducted Kyoto Encyclopedia of Genes and Genomes (KEGG) pathway enrichment analyses. Among the top 10 enriched KEGG terms, metabolic pathway, PPAR signaling, cancer pathways, cAMP signaling, and necroptosis were particularly relevant (Figure 3C). KEGG analysis revealed upregulated glycolysis/ gluconeogenesis, reduced synthesis and degradation of ketone bodies, and reduced arachidonic acid metabolism. In addition, MAPK and cAMP signaling were upregulated, while Wnt signaling was downregulated. Pathways related to cell growth and death, including cell cycle progression, necroptosis, and p53 signaling, were also enriched. In PPAR signaling pathway, ketogenesis and fatty acid oxidation were markedly downregulated, whereas lipogenesis was upregulated.

Volcano plots analysis identified 316 upregulated and 208 downregulated genes with I Log2 fold changeI ≥ 0.3 and raw *p*-value < 0.0001 were represented as (Figure 3D). The top 10 upregulated genes included *Xist*, *Gm3375*, *Dio2*, *Irf4*, *Myh6*, *Gm8126*, *Gm15543*, *Gm9732*, *Mfsd2a*, and *Mup15*, while the top 10 downregulated genes included *Il11ra2*, *Gm21451*, *Gm16867*, *Hmgcs2*, *Mid1*, *Piwil2*, *Prg4*, *Htr2b*, *Gstp3*, and *Aspg*. Among the upregulated genes, *Dio2*, which promotes brown adipocyte differentiation and thermogenesis, and *Irf4*, a key transcriptional regulator of BAT thermogenesis, were significantly upregulated. Among the downregulated genes, *Hmgcs2*, a rate-limiting enzyme of ketogenesis, was markedly downregulated. Serotonin receptor genes exhibited divergent regulation: *Htr2a* was upregulated (fold change = +2.61), while *Htr2b* was strongly downregulated (fold change = −10.46). Although serotonin regulates BAT futile cycling and thermogenesis through *Htr3*, the precise roles of *Htr2a* and *Htr2b* in BAT energy metabolism remain unclear. In addition, *Gstp3*, a regulator of glutathione transferase activity with antioxidant function, was downregulated.

### Thermoneutrality enhances survival in Ucp1-Cre;Ift88^flox/flox^ mice

Homeothermic animals must maintain their body temperatures within a narrow range to survive. However, small mammals are particularly vulnerable to heat loss at subthermoneutral conditions^15^. To test whether thermoneutral housing could rescue neonatal lethality in *Ucp1-Cre;Ift88^flox/flox^* mice with defective BAT cilia, pregnant females were housed at 30°C to provide a thermoneutral environment. After birth, littermates were reared under thermoneutral conditions, and offspring were separated from their mothers at 4 weeks of age, and male and female littermates were bred separately while being maintained at 30°C. Fourteen pups (7 *Ucp1-Cre;Ift88^flox/flox^* and 7 *Ucp1-Cre;Ift88^+/+^* mice) were maintained at 30°C. Remarkably, all *Ucp1-Cre;Ift88^flox/flox^* pups survived. At 4 weeks, body weight did not differ between *Ucp1-Cre;Ift88^flox/flox^* and *Ucp1-Cre;Ift88^+/+^* mice (Figure 4A). Histological analysis revealed smaller lipid vacuoles in BAT of *Ucp1-Cre;Ift88^flox/flox^* mice compared to *Ucp1-Cre;Ift88^+/+^* mice (Figure 4B). BAT of 4-week-old *Ucp1-Cre;Ift88^flox/flox^* mice housed under thermoneutral condition exhibited markedly increased Hmgcs2 expression relative to controls (Figure 4C), possibly reflecting a chronic compensatory response to metabolic reprogramming. Notably, *Hmgcs2* was one of the most significantly downregulated genes in P0 *Ucp1-Cre;Ift88^flox/flox^* BAT.

**Figure 4.**
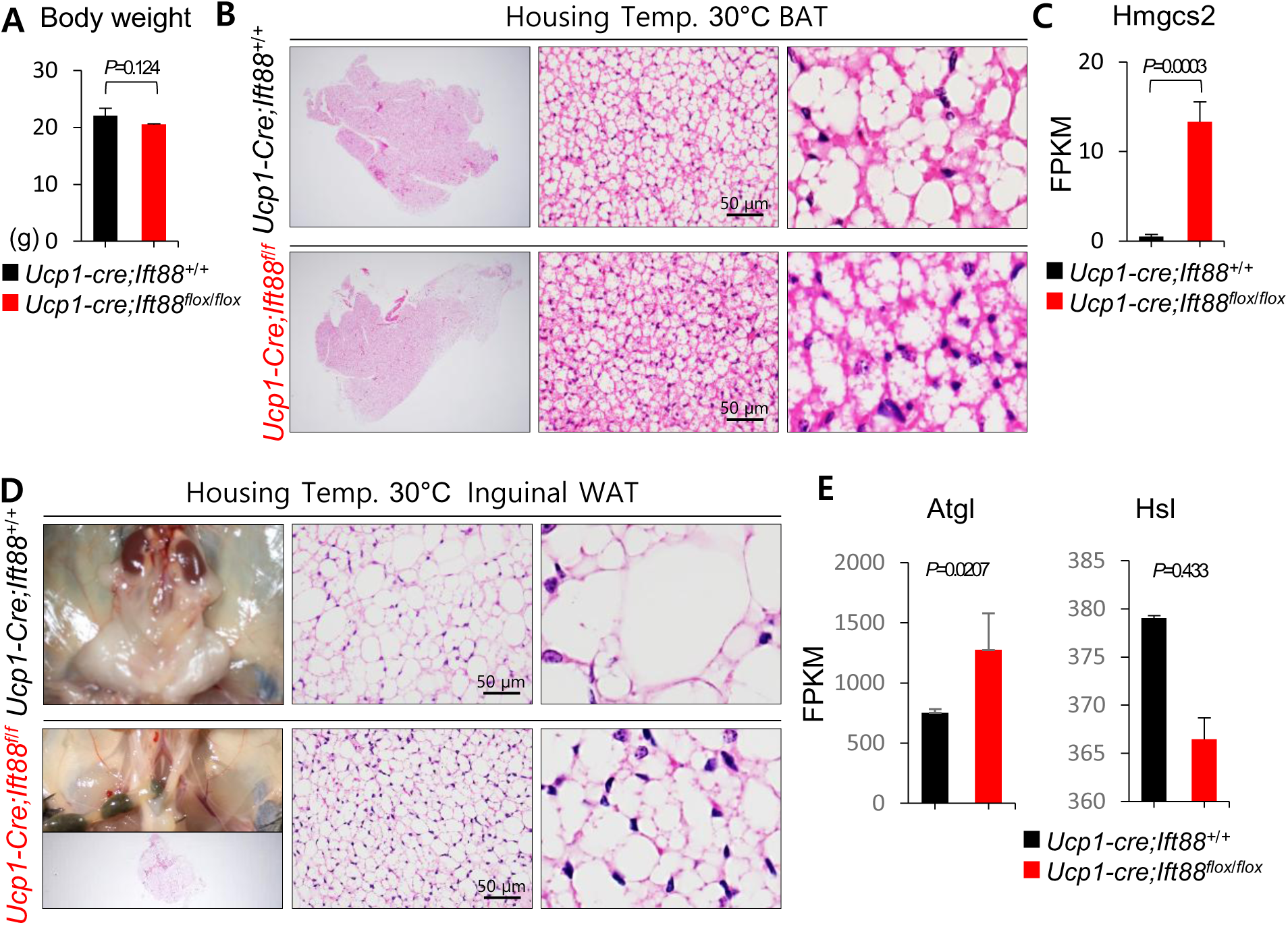
Effect of housing *Ucp1-Cre; Ift88^flox/flox^* mice under 30°C thermoneutral conditions. (A) Body weight is comparable between 4-week-old *Ucp1-Cre;Ift88^flox/flox^* and *Ucp1-Cre;Ift88^+/+^* mice. (B) In 4-week-old *Ucp1-Cre;Ift88^flox/flox^* mice, BAT lipid vacuoles are smaller than those in *Ucp1-Cre;Ift88^+/+^* mice. (C) *Hmgcs2* mRNA levels are markedly increased in BAT of 4-week-old *Ucp1-Cre; Ift88^flox/flox^* mice compared to controls. (D) WAT mass and adipocyte size are smaller in *Ucp1-Cre; Ift88^flox/flox^* mice compared to *Ucp1-Cre;Ift88^+/+^* mice. (E) *Atgl*, but not *Hsl*, expression is significantly increased in WAT of 4-week-old *Ucp1-Cre; Ift88^flox/flox^* mice.

WAT also displayed adaptations. Both the overall WAT size and lipid droplet size were significantly reduced in 4-week-old *Ucp1-Cre;Ift88^flox/flox^* mice compared to controls (Figure 4D). When BAT thermogenesis is impaired, systemic energy expenditure is disrupted, necessitating compensatory metabolic responses in WAT^16^. Consistent with this, lipolytic enzymes were examined, and *Atgl*, but not *Hsl*, was statistically significantly upregulated in *Ucp1-Cre;Ift88^flox/flox^* WAT (Figure 4E). This likely promotes fatty acid release from WAT for oxidation in other tissues to sustain energy homeostasis^17^.

Taken together, these results indicate that maintaining a suitable body temperature during the perinatal period may overcome the detrimental effects of defective primary cilia in BAT.

### Functional assessment of UCP1-dependent thermogenesis in P0 Ucp1-Cre;Ift88^flox/flox^ mice

P0 *Ucp1-Cre;Ift88^flox/flox^* mice exhibited severe hypothermia (Figure 1F), indicating a potential abnormality in thermogenesis. The expression levels of thermogenic genes-*Ucp1*, *Dio2*, *Irf4*, and *Cidea*-were comparable to or higher in the BAT of P0 *Ucp1-Cre;Ift88^flox/flox^* mice relative to controls (Figure 2B, 2C, 2E), suggesting a compensatory transcriptional response to impaired thermogenesis. To further evaluate whether UCP1-dependent thermogenesis remained functionally intact in P0 *Ucp1-Cre;Ift88^flox/flox^* mice, we performed mitochondrial respiration assays (OCR), including β3-adrenergic stimulation tests. Both OCR and extracellular acidification rate (ECAR) measurements revealed minimal differences between BAT from P0 *Ucp1-Cre;Ift88^flox/flox^* and P0 *Ucp1-Cre;Ift88^+/+^* mice (Figure 5A-5B). Treatment with β3-adrenergic agonist CL-316243 further enhanced UCP1 protein expression in BAT of both groups compared to saline-injected controls (Figure 5C).

**Figure 5.**
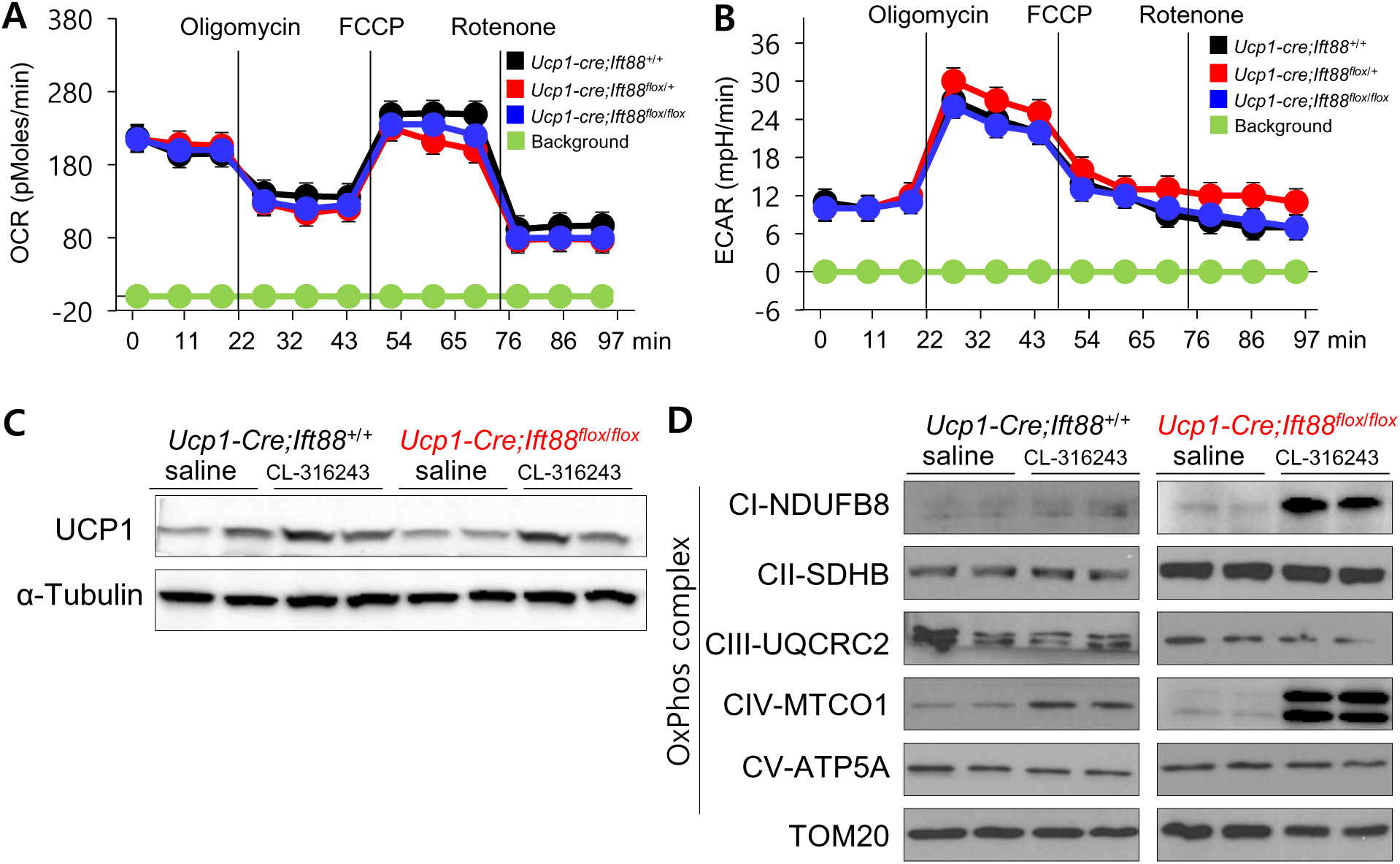
Functional assessment of Ucp1-dependent thermogenesis in P0 *Ucp1-Cre;Ift88 ^flox/flox^* mice. (A, B) OCR and ECAR in BAT show no significant differences between P0 *Ucp1-Cre;Ift88^flox/flox^* and *Ucp1-Cre;Ift88^+/+^* mice. (C) Treatment with β3-adrenergic agonist CL-316243 enhances UCP1 protein expression in BAT of both genotypes compared to saline-treated controls. (D) CL-316243 upregulates mitochondrial OxPhos complexes I and IV, while OxPhos complexes II, III, and V remained unchanged in both groups.

To assess mitochondrial thermogenic capacity more directly, we analyzed the expression of mitochondrial oxidative phosphorylation (OxPhos) complex I-V, which contribute to heat production in BAT^18^. CL-316243 administration increased the expression of complex I and IV, whereas complexes II, III, and V remained unchanged in both groups (Figure 5D). Collectively, these results indicate that, despite the severe hypothermia, the essential components of the UCP1-dependent thermogenic machinery are preserved in the BAT of P0 *Ucp1-Cre;Ift88^+/+^* mice.

### Loss of primary cilia impairs thermogenesis through mTORC1–Hmgcs2 signaling in murine BAT

Because BAT also possesses UCP1-independent thermogenic mechanisms that contribute to systemic energy and glucose homeostasis, we next investigated whether these mechanisms were preserved. In P0 *Ucp1-Cre;Ift88^flox/flox^* mice, BAT exhibited metabolic reprogramming characterized by upregulation of mitochondrial OxPhos, mTORC1 signaling, glycolysis, adipogenesis, and fatty acid metabolism (Figure 3B). These alterations likely represent compensatory mechanisms attempting to counteract defective thermogenesis. RNA-seq analysis of BAT revealed that the knockdown of *Ift88* or loss of primary cilia markedly increased the expression of genes associated with triglyceride synthesis (*Lpin1*, *Dgat1*) and fatty acid synthesis (*Acaca, Fasn, Scd1, Acly*)(Figure 6A-6B). Consistent with this metabolic shift, P0 *Ucp1*-*Cre;Ift88^flox/flox^* mice displayed hypoglycemia (Figure 1G), accompanied by upregulation of glycolysis-related genes such as *Slc2a1, Slc2a4, Pk,* and *Ldha* (Figure 6B), suggesting a compensatory adaptation to limited glucose availability.

**Figure 6.**
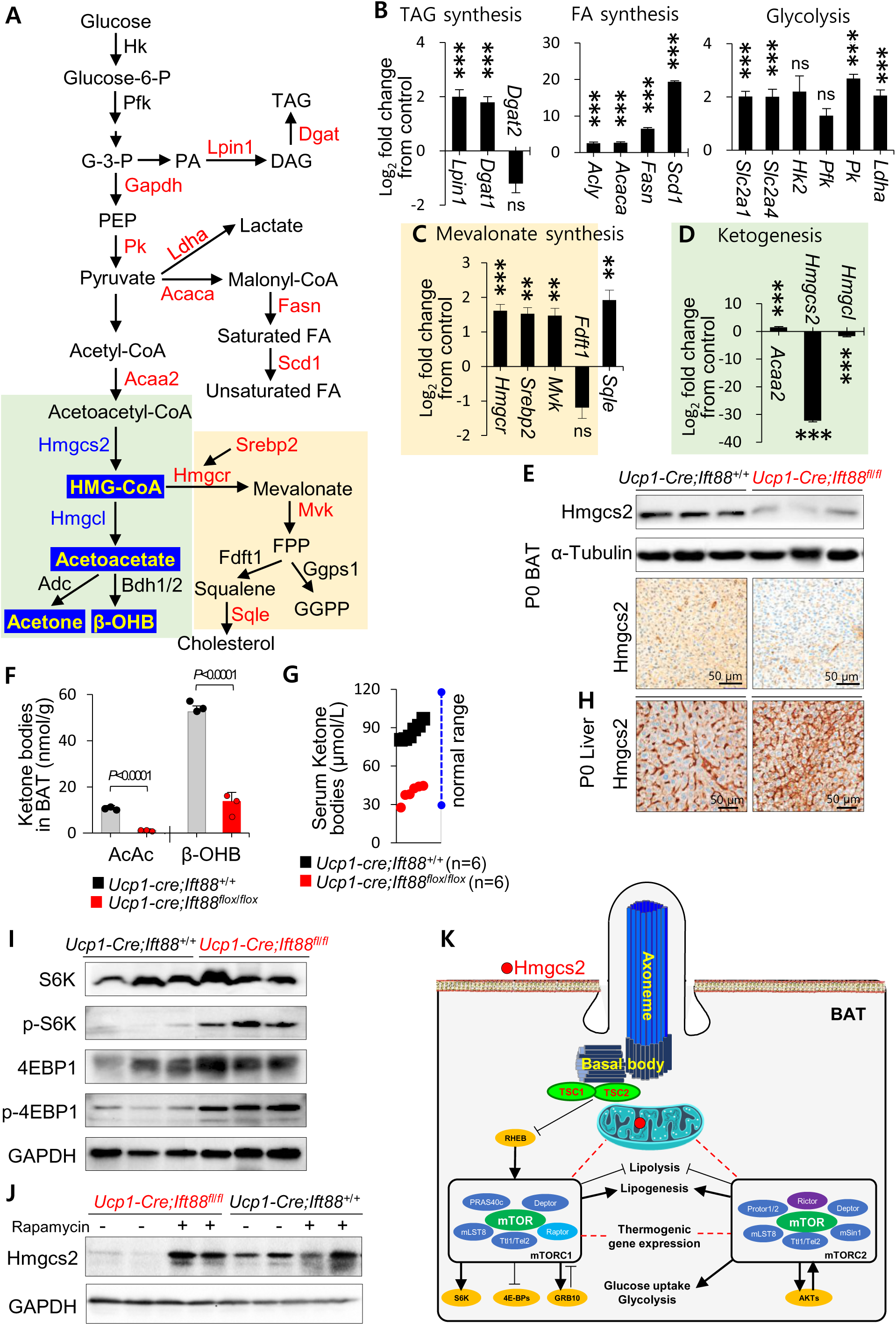
Loss of primary cilia impairs thermogenesis through mTORC1–Hmgcs2 signaling in murine BAT. (A) Diagram of triglyceride, fatty acid, glycolysis, mevalonate and ketone body synthesis pathway. (B, C) According RNA sequencing analyses of BAT, *Ift88* knockdown/ LOF of primary cilia in *Ucp1-Cre; Ift88^flox/flox^* mice significantly increase mRNA levels of genes associated with triglyceride synthesis, fatty acid synthesis, glycolysis and the mevalonate pathway. (D) mRNA expression of *Hmgcs2* and *Hmgcl*, which are required for ketone body synthesis, is most significantly reduced in *Ucp1-Cre;Ift88^flox/flox^* BAT compared to *Ucp1-Cre;Ift88^+/+^* controls. (E) Immunoblotting and IHC show decreased expressions of Hmgcs2 in *Ucp1-Cre;Ift88^flox/flox^* BAT compared to *Ucp1-Cre;Ift88^+/+^* controls. (F) BAT ketone body concentrations are quantified by GC-MS. Levels of acetoacetate and β-OHB are significantly lower in *Ucp1-Cre;Ift88^flox/flox^* mice (1.24 ± 0.57 nmol/g and 13.65 ± 4.03 nmol/g, respectively) than in P0 *Ucp1-Cre;Ift88^+/+^* mice (11.00 ± 0.97 nmol/g and 52.52 ± 2.54 nmol/g, respectively). (G) Serum levels of ketone body are decreased in P0 *Ucp1-Cre;Ift88^flox/flox^* mice (39.0 ± 6.37 μmol/L) compared to P0 *Ucp1-Cre;Ift88^+/+^* mice (86.9 ± 6.58 μmol/L). (H) Hmgcs2 IHC shows weak staining in the liver of both genotypes, with slightly increased intensity in P0 *Ucp1-Cre;Ift88^flox/flox^* mice compared to P0 *Ucp1-Cre;Ift88+/+* mice. (I) Loss of primary cilia in BAT of P0 *Ucp1-Cre;Ift88^flox/flox^* mice results in increased phosphorylation of S6K and 4EBP1 compared to controls. (J) Rapamycin treatment increases Hmgcs2 mRNA and protein expression in BAT of *Ucp1-Cre;Ift88^flox/flox^* mice and improved survival. (K) Schematic illustration summarizing that loss of primary cilia in BAT leads to aberrant mTORC1 activation, resulting in downregulation of Hmgcs2. **Abbreviation:** Hk, hexokinase; Pfk, phosphofructokinase; G-3-P, glyceraldehyde 3-phosphate; Gapdh, glyceraldehyde-3-phosphate dehydrogenase; PEP, phosphoenolpyruvate; Pk, pyruvate kinase; Acaa2, acetyl-CoA acyltransferase 2; HMG-CoA, 3-hydroxy-3-methylglutaryl coenzyme A; Hmgcl, 3-hydroxy-3-methylglutaryl-CoA lyase; Bdh1/2, 3-hydroxybutyrate dehydrogenase 1/2; β-OHB, β-hydroxybutyrate; PA, phosphatidic acid; DAG, diacylglycerol; TAG, triglyceride; Ldha, lactate dehydrogenase A; Acaca, acetyl-CoA carboxylase alpha; Fasn, fatty acids synthase; Scd1, stearoyl-Coenzyme A desaturase 1; Hmgcr, 3-hydroxy-3-methylglutaryl-CoA reductase; Srebp, sterol regulatory element binding protein; Mvk, mevalonate kinas; AcAc, acetoacetate.

We next investigated the key mechanisms required to maintain neonatal body temperature immediately after birth. The mevalonate pathway, which is essential for development and survival of brown adipocytes, is known to regulate thermogenesis^19^. However, disruption of this pathway was not observed in *Ucp1-Cre;Ift88^flox/flox^* BAT (Figure 6C). The neonatal period is characterized by a transition from the glucose-rich intrauterine environment to the postnatal stage, during which there may be a delay before regular feeding is established. During this time, neonates must rely on alternative energy sources, particularly given their limited glycogen stores. Ketone bodies serve as essential alternative energy substrates for neonates^20^. Notably, mRNA levels of *Hmgcs2* and *Hmgcl*, key enzymes in ketogenesis, were significantly decrease in P0 *Ucp1-Cre;Ift88^flox/flox^* BAT (Figure 6D). This reduction was corroborated by decreased Hmgcs2 protein expression and immunostaining intensity (Figure 6E). Consistently, the levels of acetoacetate and β-OHB were significantly reduced in BAT (Figure 6F), and serum ketone concentrations were markedly lower in P0 *Ucp1-Cre;Ift88^flox/flox^* mice (39.0 ± 6.37 μmol/L) than in P0 *Ucp1-Cre;Ift88^+/+^* mice (86.9 ± 6.58 μmol/L; normal range, 28-120 μmol/L)(Figure 6G). These findings indicate that BAT-specific ciliary loss impairs ketogenesis.

Interestingly, Hmgcs2 expression was slightly elevated in the liver of P0 *Ucp1-Cre;Ift88^flox/flox^* mice (Figure 6H). Although hepatic mitochondria are the primary site of ketogenesis, ketone body production in BAT plays a crucial role in perinatal thermogenesis^21^. Therefore, the reduced serum ketone body levels observed in P0 *Ucp1-Cre;Ift88^flox/flox^* mice are attributable to impaired BAT-derived ketogenesis. To test whether ketone body depletion accounted for neonatal lethality, β-OHB was administered intraperitoneally immediately after birth. All injected pups (2 *Ucp1-Cre;Ift88^flox/flox^*, 3 *Ucp1-Cre;Ift88^flox/+^*, 2 *Ucp1-Cre;Ift88^+/+^*) survived (Supplementary video 2, three-day-old *Ucp1-Cre;Ift88 floxed* mice).

mTOR signaling regulates diverse aspects of BAT physiology, including thermogenesis and glucose metabolism^22^. mTORC1 and mTORC2 control thermogenesis through distinct mechanisms, with each complex exerting unique and indispensable roles^22^. The TSC1/TSC2 heterodimer negatively regulates mTOR activity, and notably, TSC1 localizes to the basal body of primary cilia, suggesting that dysfunction of the primary cilium may impair TSC1/TSC2 activity and thereby promote aberrant mTOR signaling^23^. Consistent with this, loss-of-function of primary cilia in BAT from P0 *Ucp1-Cre;Ift88^flox/flox^* mice exhibited elevated mTORC1 activity, as indicated by increased phosphorylation of S6K and 4EBP1 compared to controls (Figure 6I), corroborated by RNA-seq data (Figure 3B). Pharmacological inhibition of mTORC1 with rapamycin restored *Hmgcs2* gene and protein expression in BAT and improved survival in P0 *Ucp1-Cre;Ift88^flox/flox^* neonates (Figure 6J).

Taken together, these results demonstrate that loss of primary cilia in BAT induces aberrant mTORC1 activation, leading to downregulation of *Hmgcs2* and impaired ketogenesis. These results identify a critical role for the primary cilia–mTOR signaling axis in regulating ketogenesis-related gene expression and neonatal thermogenesis in BAT (Figure 6K).

### Loss of primary cilia leads to enhanced ROS production in brown preadipocytes

BAT thermogenesis demands substantial energy. In *Ucp1-Cre;Ift88^flox/flox^* mice, both the liver and BAT fail to produce ketone bodies during the prenatal period, forcing reliance on glucose as the primary energy source. However, glucose availability significantly declines after birth. Consequently, neonatal *Ucp1-Cre;Ift88^flox/flox^* mice cannot supply sufficient fuel to BAT, resulting in impaired heat production and defective thermogenesis.

To further explore the link between defective ketogenesis and impaired perinatal thermogenesis, we conducted a series of *in vitro* experiments using murine immortalized brown preadipocytes (iBPAs) with knockdown of *Ift88* or *Kif3a*, both key genes required for primary cilium biogenesis. The knockdown efficiency of each RNA interference was confirmed at the mRNA level by RT-qPCR (Figure 7A). The siRNA-mediated knockdown of *Kif3a* (si*Kif3a*) or *Ift88* (si*Ift88*) led to the loss of primary cilia (Figure 7B).

**Figure 7.**
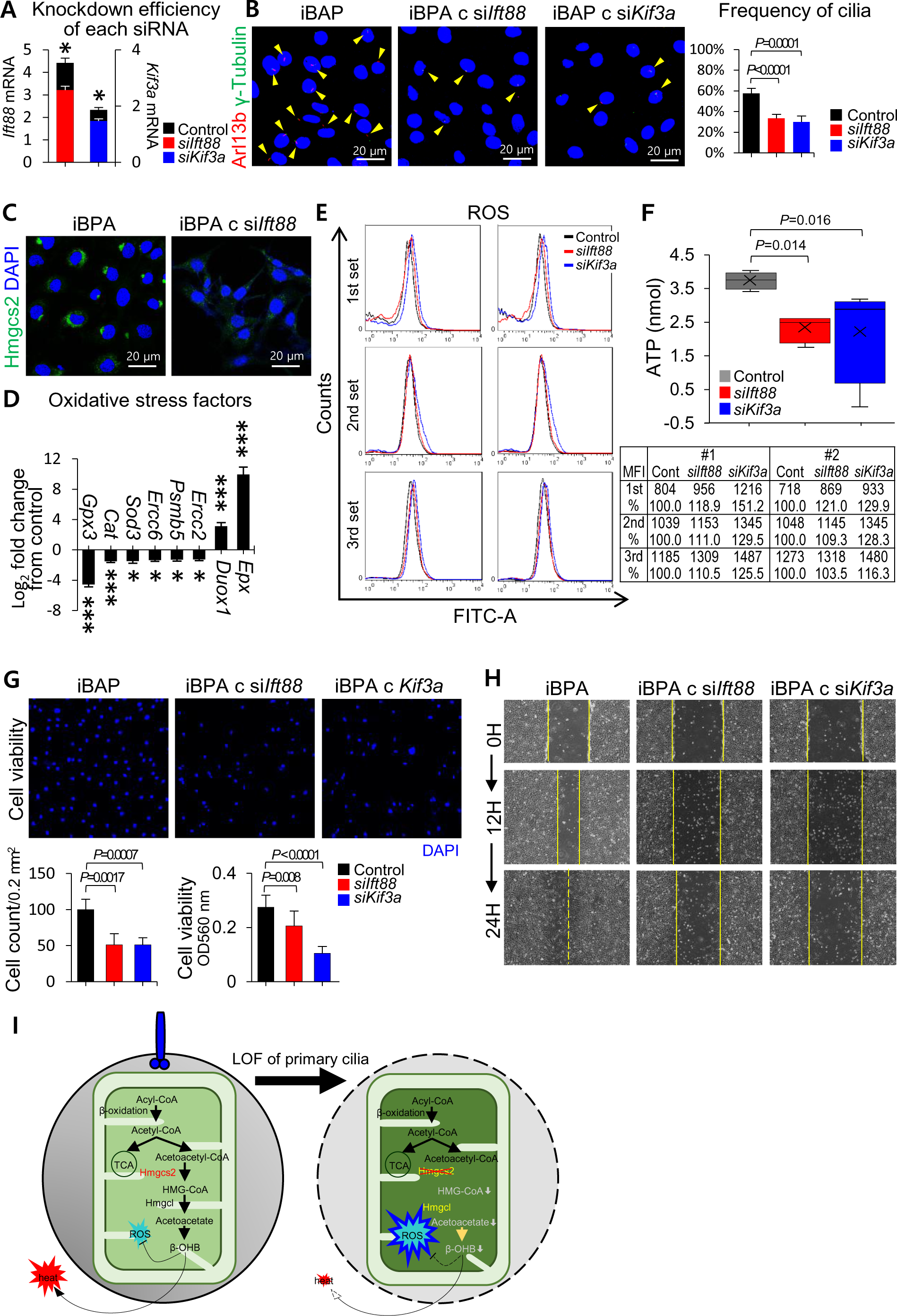
Loss of primary cilia leads to enhanced ROS production in brown preadipocytes. (A) In the murine *i*BPAs with si*Ift88* or si*Kif3a*, knockdown efficiency of *Ift88* or *Kif3a* is confirmed at the mRNA levels by RT-qPCR. *, *p < 0.05.* (B) Primary cilia are determined by IF using anti-Arl13b (for axoneme) and anti-γ-tubulin (for basal body). Primary cilia are less frequently detected in murine *i*BPA with knockdown of *Ift88* or *Kif3a.* (C) Mitochondria-localized HMGCS2 expression is decreased in *i*BPAs with ciliary loss. (D) qRT–PCR analysis of antioxidative stress factors (*Gpx3*, *Catalase*, *Sod3*, *Ercc6*, *Psmb5*, and *Ercc2*) and oxidative stress factors (*Epx* and *Duox1*) in BAT from *Ucp1-Cre;Ift88^flox/flox^ versus Ucp1-Cre;Ift88^+/+^* mice. Loss of primary cilia in BAT reduces expression of antioxidative stress factors and increases expression of oxidative stress factors in vivo. ns, not significant; *, *p* < 0.05; **, *p* < 0.005; ***, *p* < 0.001. (E) Murine *i*BPAs with *Ift88* or *Kif3a* knockdown show significantly increased H2DCFDA staining, indicating elevated ROS production. (F) Knockdown of *Ift88* or *Kif3a* reduces ATP production in murine *i*BPAs. (G, H) Knockdown of *Ift88* or *Kif3a* decreases cell viability and migration in murine *i*BPAs. (I) Schematic illustration showing that ciliary loss downregulates the ketogenic pathway through Hmgcs2 depletion, promoting ROS overproduction and subsequent BAT damage.

In line with our *in vivo* findings showing reduced *Hmgcs2* expression in the BAT of *Ucp1-Cre;Ift88^flox/flox^* mice (Figure 6E), knockdown of *Ift88* or *Kif3a* similarly reduced mitochondrial Hmgcs2 expression in murine *i*BPAs (Figure 7C).

Ketone bodies, particularly β-hydroxybutyrate (β-OHB), function not only as alternative energy substrates but also as signaling molecules with antioxidant properties^24^. Their deficiency may exacerbate oxidative stress by reducing cellular free radical scavenging capacity. We therefore investigated whether LOF of primary cilia increases mitochondrial ROS production. In the BAT of P0 *Ucp1-Cre;Ift88^flox/flox^* mice, expression of ROS-detoxification genes (e.g., *Gpx3*, *Catalase*, *Sod3*, *Ercc6*, *Psmb5*, and *Ercc2*) was suppressed, while ROS-generating genes (e.g., *Epx* and *Duox1*) were upregulated (Figure 7D). Consistently, *i*BPAs with *Ift88 or Kif3a* knockdown displayed elevated ROS levels, as assessed by H2DCFDA staining (Figure 7E).

Moreover, knockdown of *Ift88* or *Kif3a* in murine *i*BPAs reduced ATP production (Figure 7F), decreased cell viability/proliferation (Figure 7G), and impaired cell migration (Figure 7H). Together, these findings suggest that loss of primary cilia suppress *Hmgcs2* expression, leading to excessive ROS generation and increases susceptibility to cell death.

β-OHB is known to suppress ROS accumulation either directly, through antioxidant activity, or indirectly, via histone deacetylase (HDAC) inhibition and subsequent upregulation of ROS-scavenging genes^25^. Thus, ciliary loss–mediated downregulation of *Hmgcs2* impairs ketogenesis in *i*BPAs, promoting ROS overproduction, mitochondrial dysfunction, and thermogenic failure in BAT (Figure 7I).

### Loss of primary cilia in brown adipocytes does not alter fatty acid composition

Adaptive changes in the metabolic activity of BAT are generally associated with modifications in membrane phospholipid fatty acid composition^26^. We analyzed the phospholipid fatty acid composition of BAT from P0 *Ucp1-Cre;Ift88^flox/flox^* mice and their P0 *Ucp1-Cre;Ift88^+/+^* counterparts using a capillary gas chromatograph. No significant alterations were observed in the concentrations of monounsaturated or polyunsaturated fatty acids, nor in the ratios of polyunsaturated to saturated fatty acids or monounsaturated to saturated fatty acids (data not shown). The phospholipid fatty acids of BAT include C14:0 myristic acid, C16:0 palmitic acid, C16:1 palmitoleic acid, C18:0 stearic acid, C18:2 (n-6) linoleic acid, C18:3 (n-3) linolenic acid, C20:1 gadoleic acid, C20:2 icosadienoic acid, C20:3 (n-6) bis-homo-γ-linolenic acid, C20:4 (n-6) arachidonic acid (ARA), C22:5 (n-6) docosapentaenoic acid, and C22:6 (n-3) docosahexaenoic acid (DHA). P0 *Ucp1-Cre;Ift88^flox/flox^* mice showed a significantly higher level of arachidonic acid in the BAT plasma membrane compared to P0 *Ucp1-Cre;Ift88^+/+^* mice (Figure 8A). Omega-3 polyunsaturated fatty acids, including eicosapentaenoic acid (EPA) and DHA, have been shown to promote the thermogenic capacity of BAT by increasing UCP1. In contrast, omega-6 fatty acids, such as ARA, have been reported to inhibit adipocyte browning^27^. The ARA-derived n-2 prostaglandin series, including prostaglandin F receptor (*Ptgfr*) and prostaglandin E receptor 2, subtype EP2 (*Ptger2*), which are inhibitors of adaptive thermogenesis^28^, were significantly upregulated in the BAT of P0 *Ucp1-Cre;Ift88^flox/flox^* mice (Figure 8B). Specifically, mean Log2 fold changes for *Ptgfr* of P0 *Ucp1-Cre;Ift88^flox/flox^* BAT is 2.45 ± 0.45 compared to P0 *Ucp1-Cre;Ift88^+/+^* BAT (*P* = 0.004) and for *Ptger2* of P0 *Ucp1-Cre;Ift88^flox/flo^x* BAT is 3.43 ± 0.67 compared to P0 *Ucp1-Cre;Ift88^+/+^* BAT (*P* = 0.008). Overall, the fatty acid composition in BAT was not significantly altered in P0 *Ucp1-Cre;Ift88^flox/flox^* mice, suggesting that these mice maintain fatty acid homeostasis that could potentially be used for thermogenesis in the neonatal period.

**Figure 8.**
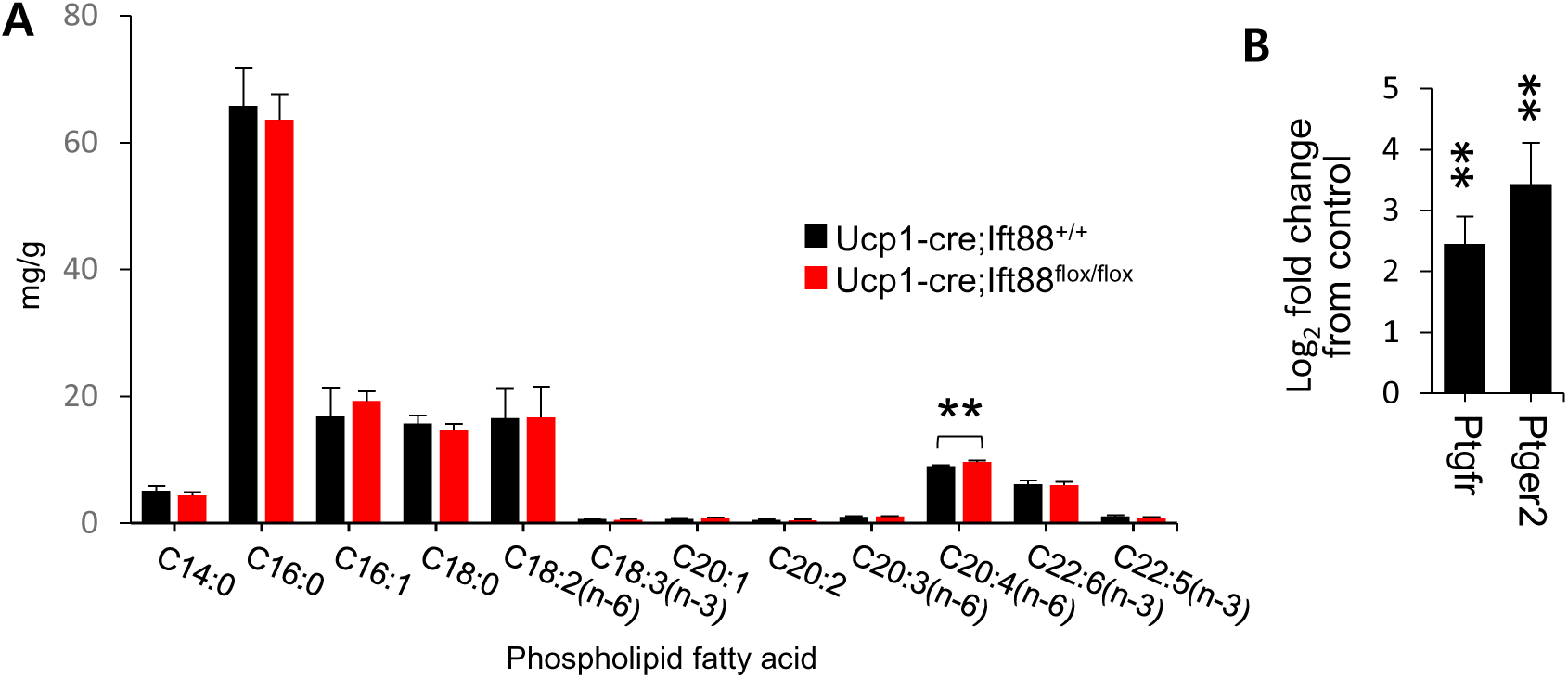
Effect of ciliary loss on phospholipid fatty acid composition of the BAT plasma membrane. (A) BAT plasma membrane from *Ucp1-Cre;Ift88^flox/flox^* mice exhibits significantly higher levels of arachidonic acid (AA) compared to *Ucp1-Cre;Ift88^+/+^* mice. Values are presented as means ± SEM for three groups (18 mice). C14:0 myristic acid, C16:0 palmitic acid, C16:1 palmitoleic acid, C18:0 stearic acid, C18:2n-6 linoleic acid, C18:3n-3 α-linolenic acid (ALA), C20:1 gadoleic acid, C20:2 eicosadienoic acid, C20:3n-6 dihomo-γ-linolenic acid (DGLA), C20:4n-6 arachidonic acid (AA), C22:5n-3 eicosapentaenoic acid (EPA), C22:6n-3 docosahexaenoic acid (DHA). (B) The ARA-derived n-2 prostaglandin series, including prostaglandin F receptor (Ptgfr) and prostaglandin E receptor 2, subtype EP2 (Ptger2), which are inhibitors of adaptive thermogenesis, are significantly upregulated in BAT from *Ucp1-Cre;Ift88^flox/flox^* mice. Mean Log2 fold changes for *Ptgfr* of *Ucp1-Cre;Ift88^flox/flox^* BAT is 2.45 ± 0.45 compared to *Ucp1-Cre;Ift88^+/+^* BAT (*P* = 0.004) and for *Ptger2* of *Ucp1-Cre;Ift88^flox/flox^* BAT is 3.43 ± 0.67 compared to *Ucp1-Cre;Ift88^+/+^* BAT (*P* = 0.008).

### Increased ciliogenesis in BAT of Ucp1-CreERT2;Ift88^+/+^ mice under cold exposure conditions

To further clarify the role of primary cilia in non-shivering thermogenesis, we examined the ciliogenesis in the BAT of adult mice exposed to chronic cold conditions. Eight-week-old *Ucp1-CreERT2;Ift88^flox/flox^* mice and *Ucp1-CreERT2;Ift88^+/+^* mice received five intraperitoneal injections of 100 µl of Tamoxifen (20 mg of Tamoxifen/1 ml of corn oil) over two weeks. Following tamoxifen treatment, mice were housed at either room temperature (23 °C) or cold conditions (6 °C) for four weeks. One of four *Ucp1-CreERT2;Ift88^flox/flox^* mice exposed to 6°C cold condition did not survive, though histological examination of its BAT revealed no obvious abnormalities.

At a room temperature (23°C), minimal histological or ciliogenesis differences were observed between BAT of *Ucp1-CreERT2;Ift88^flox/flox^* and *Ucp1-CreERT2;Ift88^+/+^* mice (Figure 9A). Under cold conditions (6°C), BAT of *Ucp1-CreERT2;Ift88^+/+^* mice displayed a relatively uniform brown adipocyte morphology with centrally located nuclei, whereas, *Ucp1-CreERT2;Ift88^flox/flox^* BAT showed heterogeneity in adipocyte size, nuclear shrinkage, and nuclear displacement to the cell periphery (Figure 9B).

**Figure 9.**
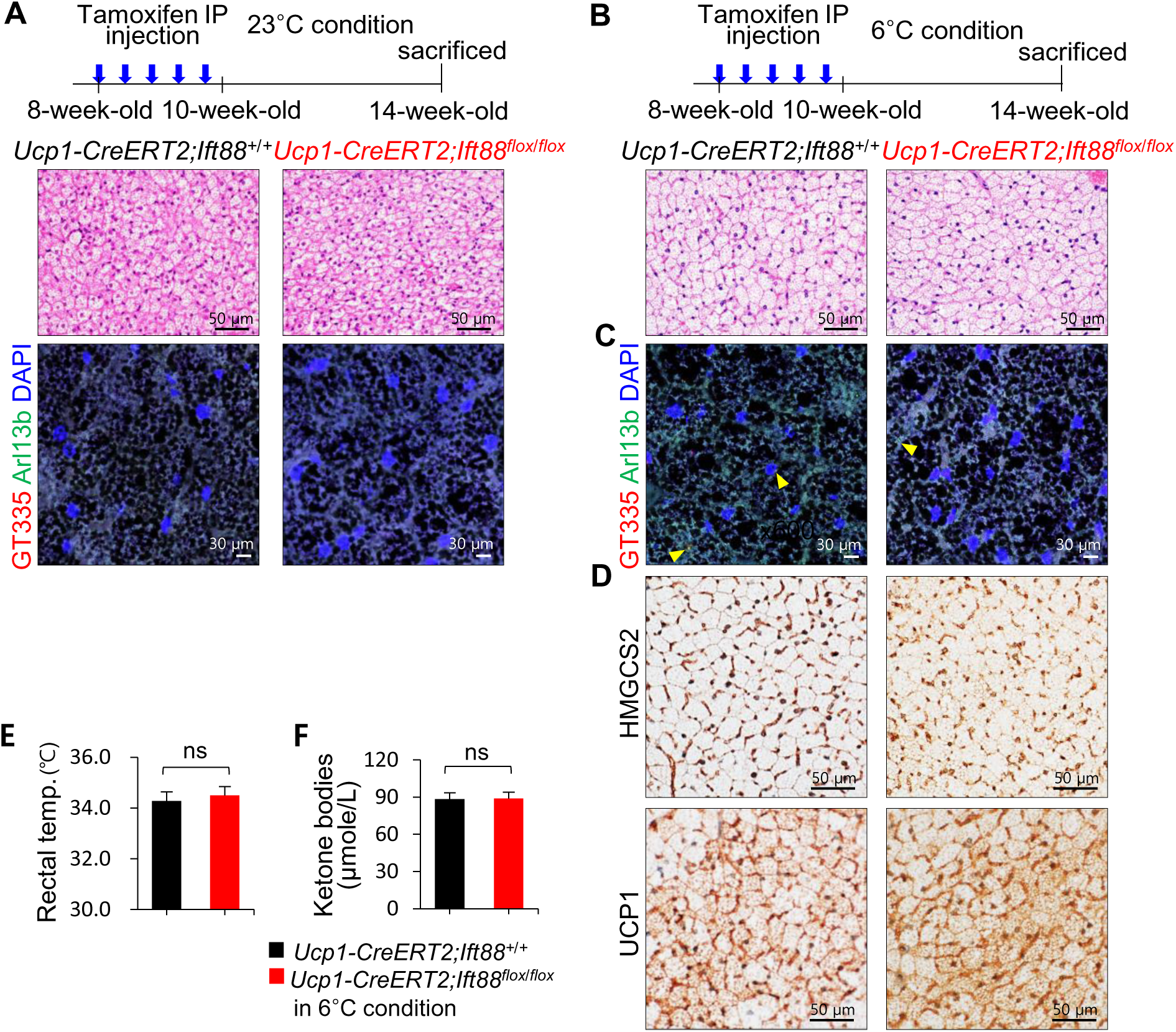
Increased ciliogenesis in the BAT of *Ucp1-CreERT2;Ift88^+/+^* mice under cold exposure. (A) At room temperature 23°C, BAT histology and ciliogenesis were similar between *Ucp1-CreERT2;Ift88^flox/flox^* mice and *Ucp1-CreERT2;Ift88^+/+^* mice. (B, C) Under cold exposure (6°C), brown adipocytes of *Ucp1-CreERT2;Ift88^+/+^* mice appears relatively uniform in size with centrally located nuclei, whereas those of *Ucp1-CreERT2;Ift88^flox/flox^* mice varies in size and shape, showing shrunken cytoplasm and peripherally displaced nuclei. IF analyses reveal an increased number of primary cilia in BAT of *Ucp1-CreERT2;Ift88^+/+^* mice (7.0 ± 1.0) compared to *Ucp1-CreERT2;Ift88^flox/flox^* mice (2.2 ± 0.8, *P* = 0.0001) exposed 6°C cold condition. (D) Rectal temperatures of *Ucp1-CreERT2;Ift88^+/+^* and *Ucp1-CreERT2;Ift88^flox/flox^* mice exposed 6°C cold condition are 34.3 ± 0.3℃ and 34.5 ± 0.3℃, respectively (*p*=0.2). (E) The levels of serum ketone body are 88.6 ± 5.0 μmol/L in *Ucp1-CreERT2;Ift88^+/+^* mice and 89.0 ± 6.4 μmol/L in *Ucp1-CreERT2;Ift88^flox/flox^* mice exposed 6°C cold condition, respectively (*p*=0.45). (F) HMGCS2 expressions are increased in BAT of *Ucp1-CreERT2;Ift88^+/+^* mice compared to *Ucp1-CreERT2;Ift88^flox/flox^* BAT exposed 6°C cold condition. ns, no significance.

Quantification revealed only a modest, non-significant reduction in primary cilia frequency in *Ucp1-CreERT2;Ift88^flox/flox^* BAT (1.2 ± 0.84) compared to controls (2.0 ± 0.70), which did not reach statistical significance (*P* = 0.07; Figure 9C). Additionally, no significant differences were detected in IHC expression of HMGCS2 and UCP1 (Figure 9D), rectal temperature (Figure 9E), or serum ketone body levels (Figure 9F). These findings suggest that primary cilia are not essential for adult BAT non-shivering thermogenesis during chronic cold exposure.

Taken together, our results indicate that while primary cilia are dispensable for cold-induced thermogenesis in mature BAT, they play a critical role in regulating ketone body metabolism during the neonatal period—a developmental stage characterized by rapid metabolic transitions and a strong reliance on ketogenesis.

## Discussion

Our findings establish that primary cilia on brown adipocytes are vital for neonatal survival under subthermoneutral conditions. In *Ucp1-Cre;Ift88^flox/flox^* mice, which lacks functional primary cilia specifically in BAT, perinatal mortality under subthermoneutral environments was complete, despite normal adipose tissue development and differentiation. While primary cilia are established as key hubs for intracellular signals during development and tissue homeostasis^29^, they appears to be dispensable for adipose development or differentiation in murine BAT. Notably, the complete rescue of perinatal lethality in *Ucp1-Cre;Ift88^flox/flox^* mice housed under thermoneutral conditions strongly suggests that primary cilia in BAT are essential for metabolic regulation during the initiation of thermogenic responses in the perinatal period.

Neonatal mice are particularly vulnerable to hypothermia because their limited white fat depots provide insufficient insulation, and their thermoregulatory centers remain immature^30^. Moreover, the natural birthing sequence often delays maternal care for several hours, prolonging neonatal exposure to ambient cold. Under these constraints, BAT-mediated non-shivering thermogenesis becomes the primary defense against hypothermia. Our data show that although the lipid composition of BAT remains largely unaltered in *Ift88* knockouts—suggesting preserved fuel stores—the mobilization or utilization of these lipids is impaired, likely due to defective ciliary regulation of key metabolic enzymes.

BAT is the principal thermogenic organ in neonatal mice and is specialized for heat production to a greater extent than skeletal muscle, particularly during the neonatal period when other thermoregulatory mechanisms are not yet fully developed^31^. In neonates, BAT primarily relies on fatty acids as its main energy source. Triglycerides stored in BAT are hydrolyzed to release free fatty acids, which are subsequently oxidized in mitochondria to generate heat through UCP1 activation^32^. Although BAT can also metabolize glucose, particularly shortly after feeding when glucose availability is high, fatty acids remain the dominant substrate for neonatal BAT thermogenesis. Our findings indicate that the fatty acid composition in BAT was not significantly altered in *Ucp1-Cre;Ift88^flox/flox^* mice, suggesting preserved fatty acid homeostasis. Interestingly, both the size of WAT depots and the lipid droplet content within WAT were markedly reduced in *Ucp1-Cre;Ift88^flox/flox^* neonates compared to controls, suggesting compensatory metabolic adaptations in WAT to maintain systemic energy homeostasis in the context of defective BAT function.

HMGCS2 is a central enzyme in ketone body biosynthesis, catalyzes conversion of acetyl-CoA into 3-hydroxy-3-methylglutaryl-CoA (HMG-CoA), a critical step in ketogenesis^33^. Ketone bodies such as β-OHB and acetoacetate are mainly synthesized in the liver, particularly during nutrient deprivation, carbohydrate restriction, or intense physical activity, and serve as alternative energy substrates, especially for the brain. In the early postnatal period, ketone bodies are particularly important because the neonatal brain expresses higher levels of ketone-metabolizing enzymes compared to the adult brain^34^. Given their limited glycogen reserves and the potential delays in regular feeding, neonates are highly susceptible to hypoglycemia. Ketone body synthesis therefore provides an additional energy supply, protecting against the deleterious effects of low blood glucose. BAT also plays an important role in non-shivering thermogenesis, and ketone bodies are thought to contribute to BAT activation, underscoring the physiological importance of Hmgcs2 and ketogenesis during early life. Arima *et al*. demonstrated that *Hmgcs2* knockdown in murine neonates reduces energy production capacity and leads to acetyl-CoA accumulation, accompanied by increased mitochondrial protein acetylation. They proposed that neonatal ketogenesis preserves mitochondria energy-producing capacity by preventing protein hyperacetylation. In line with this, the marked downregulation of Hmgcs2 observed in *Ucp1-Cre;Ift88^flox/flox^* BAT is likely a primary cause of thermogenic failure in the neonatal period. Both BAT and plasma ketone body levels were significantly reduced in *Ucp1-Cre;Ift88^flox/flox^* neonates, potentially leading to insufficient energy supply for thermogenesis and incomplete activation of UCP1, despite normal Ucp1 expression. Importantly, intraperitoneal administration of ketone bodies immediately after birth markedly improved survival, underscoring the importance of BAT-derived ketone bodies during this critical stage. Interestingly, systemic *Hmgcs2* knockdown does not produce uniformly lethal phenotypes but rather results in lethargy and increased mortality beginning after P14^34^. Although *Hmgcs2* expression is significantly reduced in *Ucp1-Cre;Ift88^flox/flox^* mice, their phenotype is distinct from systemic *Hmgcs2*-knockdown, suggesting that hypothermic death in *Ucp1-Cre;Ift88^flox/flox^* mice are as follows: no differences in blood glucose levels were observed between *Hmgcs2*-knockdown and wild type mice, while blood glucose levels were significantly lower in *Ucp1-Cre;Ift88^flox/flox^* mice. Therefore, hypothermic death in *Ucp1-Cre;Ift88^flox/flox^* neonates may reflect both *Hmgcs2* deficiency and additional consequences of ciliary loss.

Our study further demonstrates that LOF of primary cilia in BAT increases oxidative stress. Murine *i*BPAs with defective primary cilia exhibited enhanced susceptible to oxidative stress, resulting in an excessive ROS production. Ketone body metabolism is known to protect against oxidative damage by inhibiting ROS generation and enhancing endogenous antioxidant defenses^35^. Consistent with this, Nishitani *et al*. reported that Hmgcs2 regulates antioxidant factors and ROS in adipocytes via autocrine signaling though endogenous ketone body production^36^. Accordingly, ROS overproduction caused by impaired ketogenesis likely contributes to the defective thermogenesis observed in P0 *Ucp1-Cre;Ift88^flox/flox^* neonates.

We also found that mitochondria-localized Hmgcs2 levels decrease in *i*BPAs lacking primary cilia, suggesting possible communication between primary cilia and mitochondria. Although several studies have suggested crosstalk between cilia and mitochondria, the mechanisms remain poorly understood. Burkhalter *et al*. recently presented electron micrographs indicating a possible tubular connection between some mitochondria and the base of the primary cilium^37^. Consistent with this, Noah Moruzzi *et al*. demonstrated a spatial relationship between the cilium and the mitochondrial network, showing that mitochondrial impairment and ROS overproduction can also influence ciliary morphology and number^38^. The precise mechanisms by which the loss of primary cilia leads to an overall reduction in Hmgcs2 expression remain to be elucidated. Clarifying these mechanisms will provide critical insights into the pathophysiological processes underlying hypothermic death in newborn *Ucp1-Cre;Ift88^flox/flox^* mice.

In summary, this study demonstrates that primary cilia in BAT are indispensable for neonatal thermogenesis and survival under cool ambient conditions, primarily through regulation of Hmgcs2-mediated ketone body production. Loss of primary cilia disrupts ketogenesis, increases oxidative stress, and impairs non-shivering thermogenesis, leading to perinatal lethality that can be fully rescued by thermoneutral housing or ketone body supplementation. These findings reveal a critical cilium–mitochondria metabolic axis in early life and provide a framework for exploring ciliary modulation as a potential therapeutic strategy in ciliopathy-associated metabolic dysfunction.

## Acknowledgements

This work was supported by the Basic Science Research Program through the National Research Foundation of Korea (NRF), funded by the Ministry of Science, ICT (MISIT)(grant number: NRF-2020R1A2C2010269); 2023 Research Settlement Fund for the New Faculty of Konyang University Hospital.

## Author contributions

Junguee Lee, Yong Min Kim: designed research, performed research, analyzed data, wrote the paper.

Jaeku Kang: designed research, analyzed data, wrote the paper.

Junguee Lee: Funding acquisition

Seung Yun Han, Hwan-Woo Park, Jongdae Shin, Jong-seok Kim, Hyeseung Lee, Joo Heon Kim, Hyung

Seo Park: contributed new reagents or analytic tools, analyzed data.

## Supplemental Information

**Supplementary video S1**

P0 *Ucp1-Cre;Ift88^+/+^* pups actively shiver, whereas P0 *Ucp1-Cre;Ift88^flox/flox^* pups remain relatively immobile at an ambient temperature of 25°C.

**Supplementary video S2**

*Ucp1-Cre;Ift88* floxed mice weree injected intraperitoneally with β-OHB immediately after birth. The video shows the surviving three-day-old mice (2 *Ucp1-Cre;Ift88^flox/flox^* ; 3 *Ucp1-Cre;Ift88^flox/+^* ; 2 *Ucp1-Cre;Ift88^+/+^* pups).

